# Genome-Wide Covariation in SARS-CoV-2

**DOI:** 10.1101/2021.03.08.434363

**Authors:** Evan Cresswell-Clay, Vipul Periwal

## Abstract

The SARS-CoV-2 virus causing the global pandemic is a coronavirus with a genome of about 30Kbase length [Song et al., 2019]. The design of vaccines and choice of therapies depends on the structure and mutational stability of encoded proteins in the open reading frames(ORFs) of this genome. In this study, we computed, using Expectation Reflection, the genome-wide covariation of the SARS-CoV-2 genome based on an alignment of ≈ 130000 SARS-CoV-2 complete genome sequences obtained from GISAID[Shu & McCauley, 2017]. We used this covariation to compute the Direct Information between pairs of positions across the whole genome, investigating potentially important relationships within the genome, both within each encoded protein and between encoded proteins. We then computed the covariation within each clade of the virus. The covariation detected recapitulates all clade determinants and each clade exhibits distinct covarying pairs.

## 1 Introduction

Severe Acute Respiratory Syndrome (SARS) is a viral respiratory disease caused by the SARS-associated coronavirus. In December 2019, this pneumonia-like disease re-emerged in the Chinese city of Wuhan and the novel beta-coronavirus 2 (SARS-CoV-2) was identified as the causative agent [Zhu et al., 2020]. The genome was first characterized by Wu et al. in December 2019 [Wu et al., 2020]. Since then the SARS-CoV-2 virus has spread relentlessly all over the world and been declared a worldwide pandemic with 79 million cases leading to 1.7 million deaths to date [World Health Organization, 2020, John Hopkins University, 2020]. SARS-CoV-2 is an +ssRNA virus belonging to the coron-aviridae family major genera Betacoronavirus [Gorbalenya et al., 2020]. The viral genome encodes several open reading frames (ORFs): ORF1ab, ORF3a, ORF6, ORF7a, ORF7b, ORF8, ORF10. These ORFs encode for several non-structural proteins (NSPs) while there are specific regions encoding the spike glycoprotein (S), envelope (E), membrane glycoprotein (M), and the nucleocapsid protein (N). The genome (NC_045512.2, 29870 nucleotides long) of the virus can be broken into 11 encoding regions: ORF1ab (266-21555), S (21563-25384), ORF3a (25393-26220), E (26245-26472), M (26523-27191), ORF6 (27202-27387), ORF7a (27394-27759), ORF7b (27756-27887), ORF8 (27894-28259), N (8274-29533), ORF10 (29558-29674) [NCBI, 2020].

While the reference genome is used for most investigations, there is also an abundance of data available which can be used to monitor variations in the genome and analyse the the evolution and nature of the virus. This data was assembled by GISAID to document different strains of the virus in a new database: EpiCoV. With the first viral entry on January 10 2020, the database has grown to 292,000 submissions [Shu & McCauley, 2017]. In this study we use 137636 of these strains to analyse the evolution of the virus.

In our analysis we develop a co-evolutionary interaction network of nucleotide positions using an entropy-based method to infer genome-level interaction in the SARS-CoV-2 genome.

### 1.1 Coevolution

The variation of the virus’s genetic structure is of considerable medical and biological importance for prevention, diagnosis, and therapy. Mutations in the viral genome allows us to investigate potentially important relationships within the genome. Comparative RNA sequence sequence analysis has long been used to investigate co-evolution via covariance of nucleotide mutations (30,31) with difficulty arising in the separating of indirect and direct interactions that lead to such co-variation. A similar issue in inferring protein residue interactions was addressed by Direct Coupling Analysis (DCA) which successfully inferred direct interactions (DIs) in proteins as well as between proteins [Weigt et al., 2009]. Recently DCA was also applied to RNAs and RNA-protein complexes [De Leonardis et al., 2015] For RNAs, the DCA-based methods infer physical interactions, both secondary and tertiary, between nucleotides in an RNA molecule by analyzing the co-evolutionary signals of nucleotides across sequences in the RNA family [De Leonardis et al., 2015]. In this paper we will utilize Expectation Reflection [Hoang et al., 2019a, Hoang et al., 2019b] with DCA to infer DI in the SARS-CoV-2 genome. These interactions may also provide information on protein-protein interaction. Additionally this analysis could be useful in vaccine development, aiding in efforts to mitigate “escape pathways” for the virus to use in future strains [Dahirel et al., 2011].

### 1.2 SARS-CoV-2 Genome Analysis using Expectation Reflection

Our analysis begins with the acquisition and alignment of genome sequence data, described in Section 4.2. Once the data is aligned we pre-process the aligned sequences by removing sequence positions which contain 95% or more conservation as discussed in Section 2.1. We infer covarying positions from the curated-aligned genome data using DCA-ER which is outlined in Section 4.1. The resulting genome-wide interactions are discussed by encoding region in Section 2.2. Our analysis includes the presentation of position interaction maps and tabulation of the strongest resulting DI pairs. For the strongest DI pairs we also present the single site amino acid (AA) frequency as well as the AA-pair counts. Similar analysis is also applied to the G, GR, GH, S and V clades in Section 2.3.

## 2 Results

### 2.1 Clade Incidence

As in any ab initio inference problem, when inferring co-evolutionary interactions between nucleotide positions in the SARS CoV-2 genome, we must consider certain properties of the data at hand. As an example, the length of the full genome is approximately 29000 nucleotide positions. However, when we consider genome positions in which no single nucleotide is expressed more than 95% of the time (95% conserved), the relevant positions are reduced by approximately two-thirds. Decreasing the conserved percentage decreases the number of allowed repetitions in a given position (data column). In other words, as we decrease the threshold for conserved columns, the condition for variation at a given position becomes more stringent and the number of columns retained for analysis will decrease. As in the example above, going from 100% (full genome, all columns allowed) to 95% conservation removes a significant number of genome positions. Because inference with ER relies on mutations at a given position we must consider the resulting number of columns, or incidence, after such curation. In addition, region-specific incidence may also underline importance to the efficacy of the virus because higher incidence represents more variation and mutation. We also consider the clade-specific incidence of the full genome since we will consider genome interactions in different clades in future sections.

In Figure 1 we plot the incidence of the ORF1ab and S regions for different thresholds of allowed conservation. These incidence curves are given for the different clade data sets. Figure 1 shows that region incidence varies between clades and that the incidence of different encoding regions is affected differently for a given clade. For example, consider the full genome data set against the S clade set in Figure 1. The full genome data set (blue) has one of the highest ORF1ab incidence curves, but the same level of incidence is not necessarily expressed in the S region. In contrast, the S clade (purple) shows middling incidence for ORF1ab and one of the strongest incidence curves for the S encoding region.

**Figure 1:**
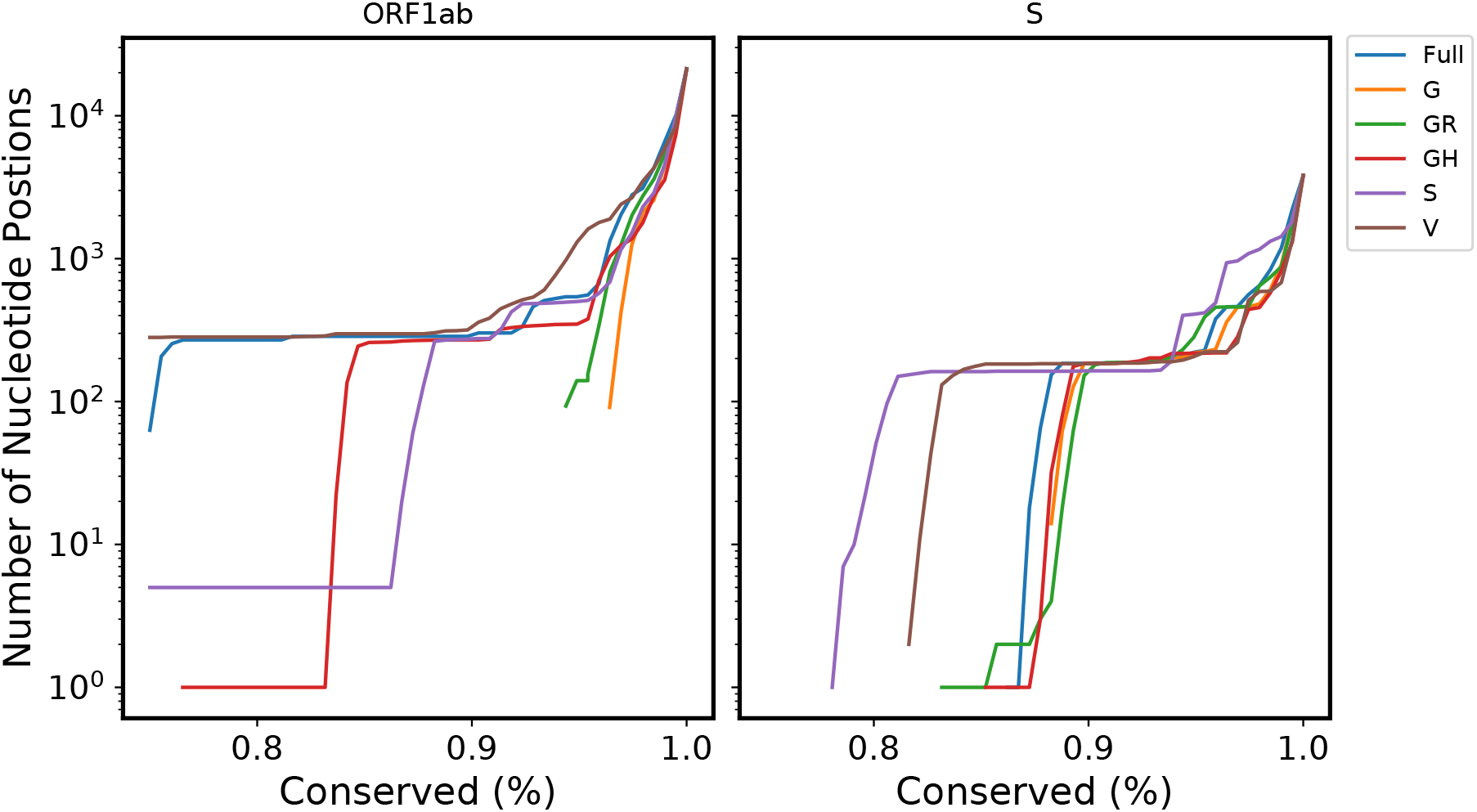
Comparing Incidence across Clades. We plot total number of retained columns (incidence) against the allowed conserved percentage per a given nucleotide position for both ORF1ab and S regions (across all clades).

This analysis can also give some intuition on the nature of individual encoding regions by quantifying the level of variability expressed within a given clade. For example, in Figure 1, we see differences in which clades express higher incidence between the ORF1ab and S encoding regions. We can apply the same principle to compare the ORF1ab and S regions in the full genome data set. In Figure 2, we plot the incidence of ORF1ab and S in the full genome sequence set as we increase the allowed conserved percentage per given nucleotide position. This figure shows different levels of position-wise variability for the two encoding regions.

**Figure 2:**
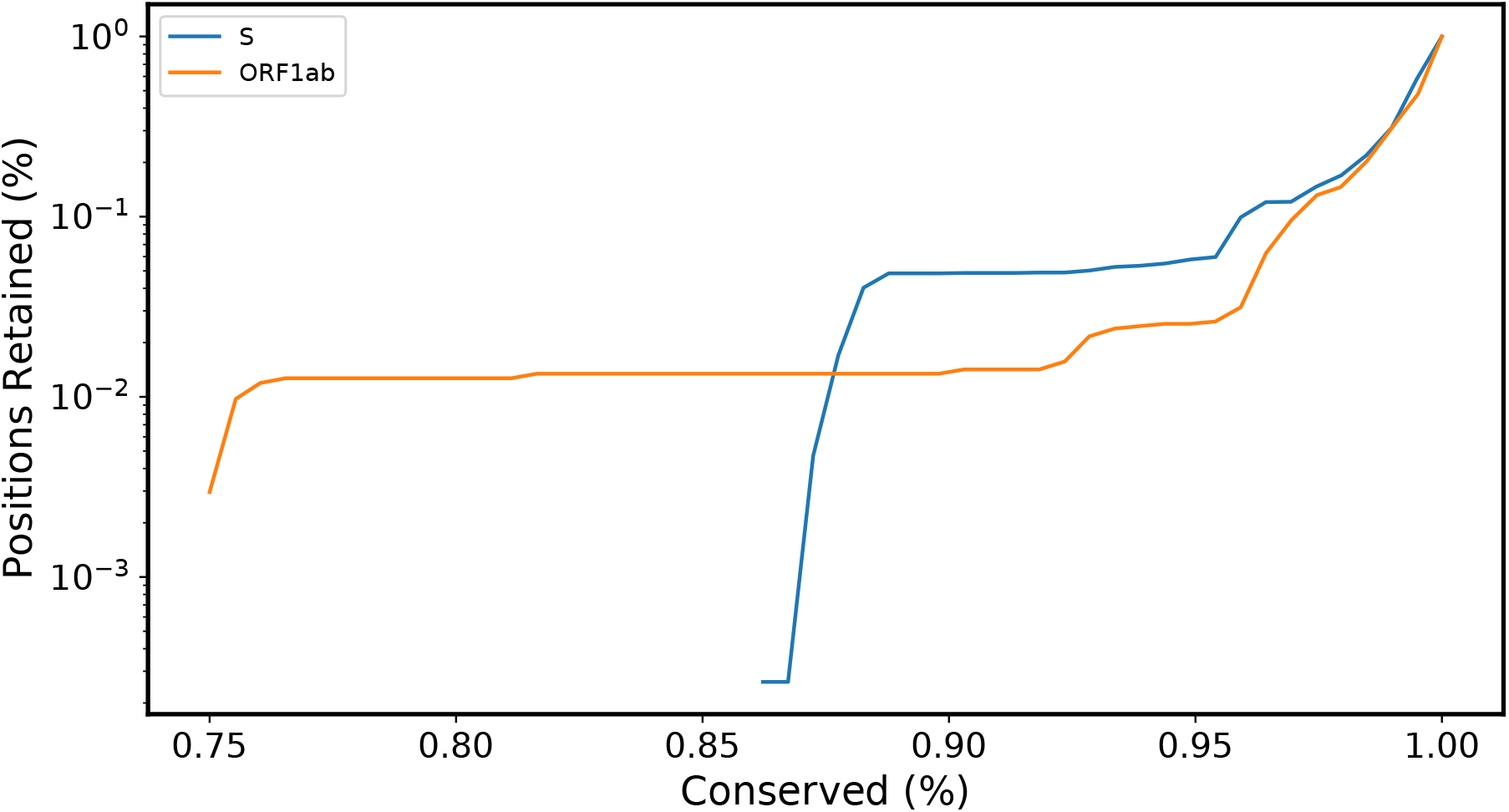
Comparing the Incidence of S and ORF1ab. We plot the percentage of retained columns (incidence) against the allowed conserved percentage per a given nucleotide position (in the full genome data set).

For the remainder of the paper we will set the conservation threshold to 95%. This is in order to retain a significant number of positions for the subsequent analysis. We must also consider that the size of a given clade, or genome data set will affect the incidence and variability. Specifically, as the number of sequence considered changes, the level of variability, enforced by the conservation thresholds, will be altered as well. Therefore, when considering smaller data sets, such as the S and V clades, we must keep in mind that the incidence is affected by the cardinality of the set itself.

### 2.2 Genome Wide Analysis

We begin by inferring interactions between nucleotide positions across the entire genome. Figure 3 shows a gray-scale of DI calculated from inferred couplings between positions (*i, j*) using all available sequences. Position pairs which showed significant coevolution (*DI* ≥ .1) are emphasized. In Figure 3, the full interaction map is dominated by interactions in ORF1ab (positions 266-21556), S (positions 21564-25384), and ORF7ab (27395-27888). Proximal nucleotide positions (diagonal of the interaction map) express strong covariation as shown by the thick black diagonal bar. In fact, we suppress the emphasis of *DI >* 0.1 for proximal pairs (∥ *i* − *j* ∥ < 10) in all interaction maps due to the prevalence of proximal pair interactions with *DI >* 0.1. However, there are several off-diagonal position pairs (far apart in the genome) which show strong covariation. This shows potential evolutionary links between specific positions or regions in different parts of the genome. As an example we can consider the top 5 DI pairs for the entire genome in Table 2.2. While most of the position pairs are proximal, the strongest interaction is more than 20000 nucleotides apart.

**Figure 3:**
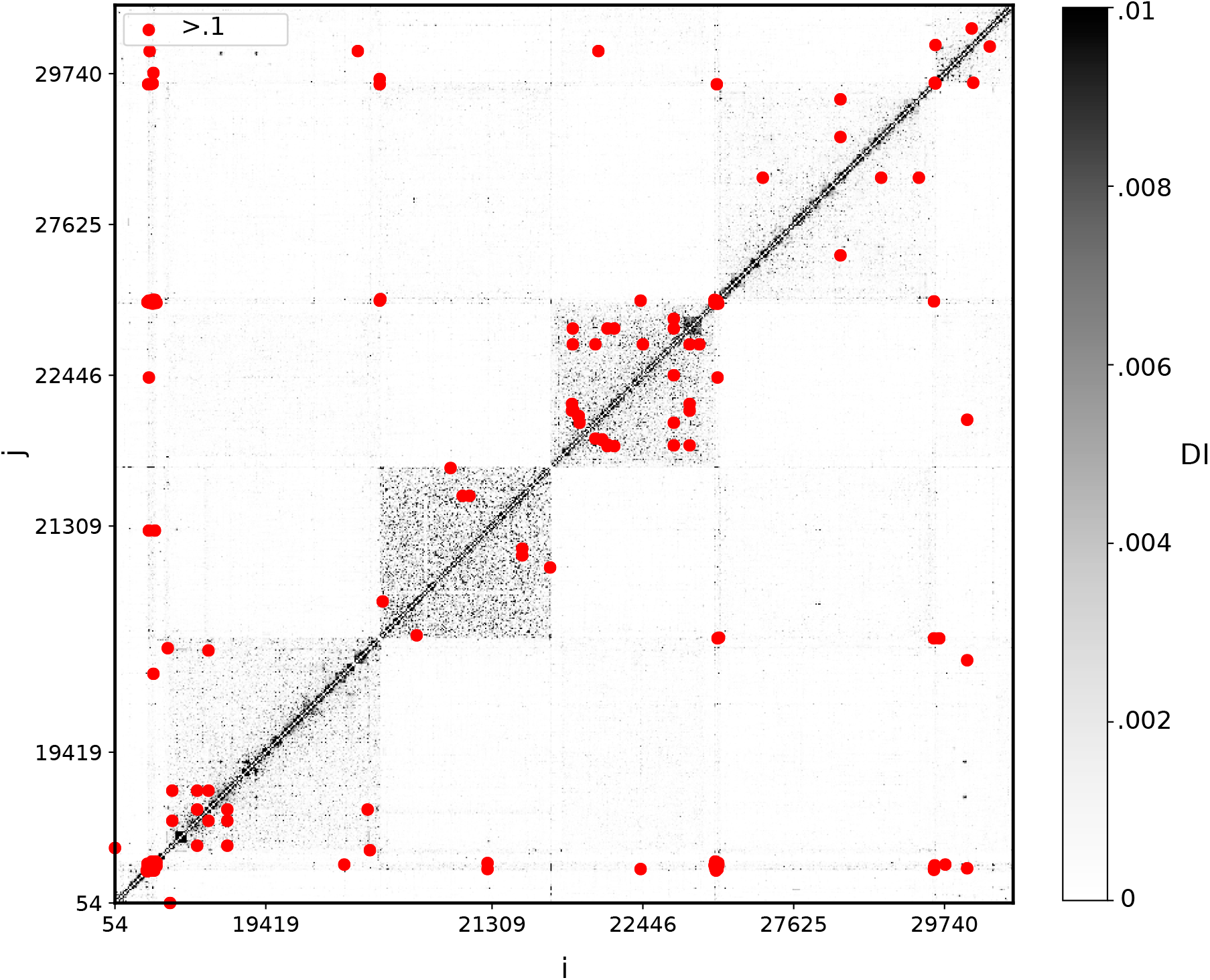
hCoV-19 Genome-Wide Interaction Map. We infer covariation in nucleotide positions across ≥ 130000 sequences.

**Figure 4:**
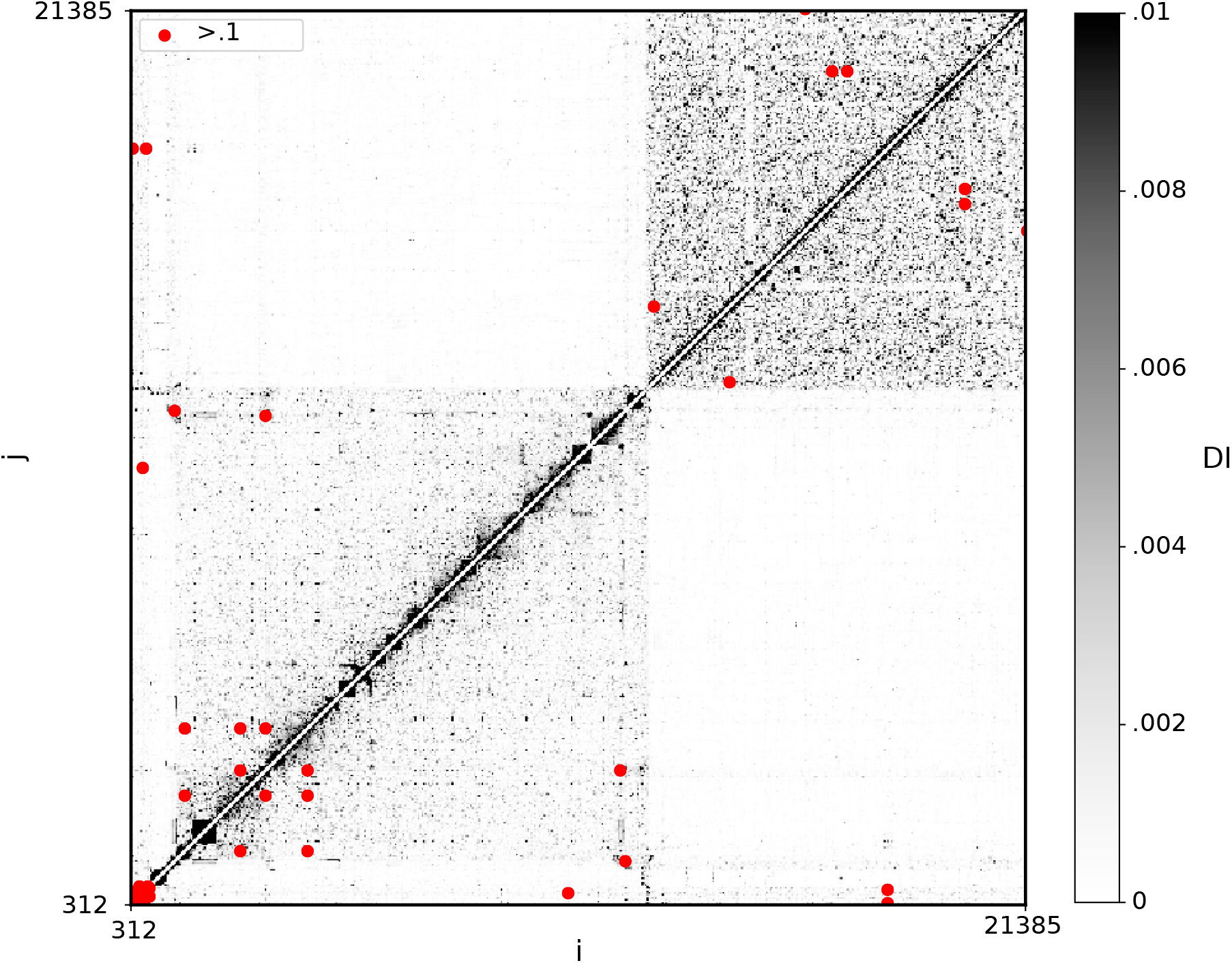
ORF1ab Interaction Map.

| Position 1 | Position 2 | DI |
| --- | --- | --- |
| 2237 | 22384 | 0.915 |
| 28881 | 28883 | 0.801 |
| 29700 | 29721 | 0.670 |
| 241 | 313 | 0.665 |
| 14408 | 3037 | 0.456 |

In order to further explore features from the full interaction map we will divide the full genome into different encoding regions, focusing on those regions which show significant incidence.

#### 2.2.1 ORF1ab

The ORF1ab region of SARS-CoV-2 genome is an important polyprotein gene which encodes 16 nonstructural proteins important to the life cycle of the virus. Because of this importance, some of the proteins encoded in this region have been proposed as potential targets for antiviral therapy [Kwong et al., 2005, Briguglio et al., 2011, Zhou et al., 2020]. Figure 3 shows that the region also has the largest number of non-conserved nucleotide positions, or position incidence, (as described in Section 2.1) of the major encoding regions. This increased incidence is expressed in the cardinality of the interaction map.

Table 1 shows the top 10 DI pairs in ORF1ab with bolded positions representing distal (non ORF1ab) positions. Note that more than half of the top 10 DI pairs in Table 1 are distal which may be an indicator of the significance of the region to other encoding regions (and their resulting functions).

**Table 1:** Top 10 ORF1ab DI pairs. Listing the strongest DI pair positions (bolded postions not in ORF1ab)

| Position 1 | Position 2 | DI |
| --- | --- | --- |
| 14408 | 3037 | 0.456 |
| 1059 | **25563** | 0.441 |
| 14805 | **26144** | 0.380 |
| 7540 | **23401** | 0.343 |
| 14408 | **23404** | 0.340 |
| 3037 | **23403** | 0.335 |
| 21254 | **22227** | 0.329 |
| 18555 | **23401** | 0.313 |
| 21367 | 21369 | 0.291 |
| 1163 | **23401** | 0.283 |

While the interaction map and DI pairs are a useful overview of genome coevolution, it is important to consider whether these coevolving positions result in alterations of amino acids encoded by the interacting positions. Table 2 gives the resulting proportion of amino acids (AA) encoded by the nucleotide positions outlined in Table 1. The table shows frequent occurrence for a dominant AA at any given position. This analysis can be extended to consider the AA pair counts for a given genome position pair. Table 3 shows the AA pair counts of the dominant DI pairs in ORF1ab. In the given position pair matrices we see three cases arise: A high AA-pair count on the diagonal, on a single row/column, or in a single element. A high AA-pair count on the diagonal means the prevalence of two AA-pairings resulting from the coevolution of the two positions. The AA-count between positions 14408 and 23404 is a good example of this first case. Dominance of a given row or column suggests that one position remains mostly fixed while the other position expresses variability, as seen in the AA-counts for the position position pairing of 3037 and 23403 where 3037 almost always encodes Phenylalanine. The final case in Table 3 is a single AA-pair expressed dominantly for a given position as seen with positions 21367 and 21369. This case shows little coevolution and generally occurs in AA-pairs from positions with lower DI where both positions have a dominant AA in their single site AA frequency (see 21367 and 21369 in Table 2). We continue with the same analysis for the S encoding region.

**Table 2:** ORF1ab Single Site AA Frequencies. We consider the distributions of resulting amino acids from the nucleotide positions which had the highest DI in ORF1ab

|  | Single Site<br>Freqs. |
| --- | --- |
| 14408 | L: .857, P: .139 |
| 3037 | F: .995 |
| 1059 | T: .832, I: .164 |
| 25563 | Q: .783, H: .215 |
| 14805 | Y: .998 |
| 26144 | G: .945, V: .040 |
| 7540 | T: .997 |
| 23401 | Q: .999 |
| 23404 | G: .861, D: .138 |
| 23403 | G: .861, D: .138 |
| 21254 | A: .933, X: .106 |
| 22227 | A: .956, V: .029 |
| 18555 | D: .999 |
| 21367 | I: .941, X: .059 |
| 21369 | I: .941, X: .059 |
| 1163 | I: .938, F: .059 |

**Table 3:** ORF1ab AA Pair Counts. Encoded amino acid counts for high ranked DI positions in ORF1ab.Considering the most prevalent amino acids for each position.

|  |  |  |  |  |  |  |  |  |  |  |
| --- | --- | --- | --- | --- | --- | --- | --- | --- | --- | --- |
| 3037<br>14408 | **F** | **other** | 14408<br>23404 | **L** | **P** | **other** | 1059<br>25563 | **T** | **I** | **other** |
| **L** | 117404 | 497 | **G** | 117540 | 457 | 521 | **Q** | 107023 | 257 | 434 |
| **P** | 19006 | 117 | **D** | 269 | 18615 | 72 | **H** | 7274 | 22237 | 99 |
| **other** | 544 | 68 | **other** | 101 | 40 | 21 | **other** | 223 | 45 | 44 |
| 3037<br>23403 | **F** | **other** | 21254<br>22227 | **A** | **X** | **other** | 14805<br>26144 | **Y** | **other** |  |
| **G** | 117983 | 535 | **A** | 123066 | 8487 | 14 | **G** | 129808 | 259 |  |
| **D** | 18827 | 129 | **V** | 3845 | 246 | 0 | **V** | 5532 | 12 |  |
| **other** | 143 | 19 | **other** | 1452 | 525 | 1 | **other** | 1988 | 37 |  |
| 21367<br>21369 | **I** | **X** | **other** | 1163<br>23401 | **I** | **F** | **other** |  |  |  |
| **I** | 129512 | 0 | 0 | **Q** | 128884 | 8055 | 488 |  |  |  |
| **X** | 0 | 8112 | 0 | **other** | 172 | 23 | 14 |  |  |  |
| **other** | 0 | 0 | 12 |  |  |  |  |  |  |  |

#### 2.2.2 Spike Glycoprotein

The Spike protein encoding region (S) plays a vital role in viral entry into the host cells [Gallagher & Buchmeier, 2001]. As a result this region is considered a key target in current vaccine development [Tai et al., 2020]. In Figure 5 and Table 4 we present both the interaction map and the top ranked DI pairings respectively for the S region gene positions. In addition to this DI analysis we present the single site frequencies and AA-pair non-singular AA-counts in Tables 5 and 6. For the S encoding region we only include the top 3 rated DI amino acid pairings. Regardless of this curation, the AA-pair matrices in Table 6 show a single AA-pair count prevalence for all but the strongest DI pairing. This pairing, between positions 22363 and 22384, shows a strong diagonal. However, the alternate can be any amino acid pairing which shows that the main trend is the threonine-valine combination.

**Table 4:** Top 10 S DI pairs. Listing the strongest DI pair positions (bolded positions not in S)

| Position 1 | Position 2 | DI |
| --- | --- | --- |
| 22363 | 22384 | 0.915 |
| 23401 | **7540** | 0.343 |
| 23401 | **18555** | 0.313 |
| 22497 | 22495 | 0.301 |
| 22353 | 22355 | 0.292 |
| 23401 | **1163** | 0.283 |
| 23401 | **16647** | 0.275 |
| 22334 | 22336 | 0.263 |
| 22539 | 22541 | 0.245 |
| 22526 | 22524 | 0.239 |

**Table 5:** S Single Site AA Frequencies. We consider the distributions of resulting amino acids from the nucleotide positions which had the highest DI in S

|  | Single Site<br>Freqs. |
| --- | --- |
| 22363 | V:.894,X:.106 |
| 22384 | T:.896,X:.104 |
| 23401 | Q:.998,X:.001 |
| 7540 | T:.997,X: .003 |
| 18555 | D:.997,X:.003 |
| 22497 | I:.893,X:.107 |
| 22495 | G:.893,X:.107 |
| 22353 | A:.891,X:.109 |
| 22355 |  |
| 1163 | I:.938,X:.057 |
| 16647 | T:.989,X:.011 |
| 22334 | W:.937,X:.062 |
| 22336 | W:.938,X:.062 |
| 22539 | I:.951,X:.049 |
| 22541 | V:.994,X:.005 |
| 22526 | P:.947,X:.053 |
| 22524 | Q:.937,X:.062 |
| 22354 | A:.891,X:.109 |
| 22488 | E:.893,X:.106 |

**Table 6:**
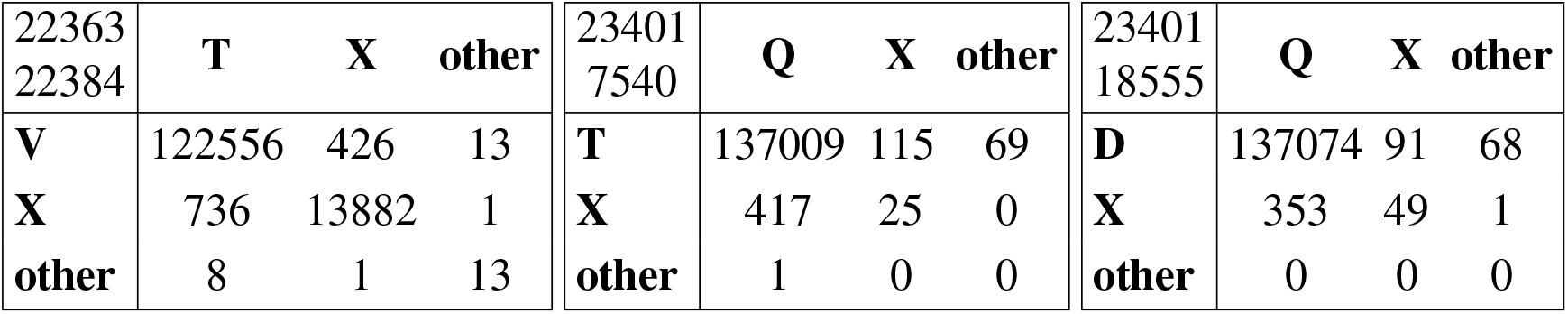
S AA Pair Counts. Encoded amino acid counts for high ranked DI positions in S. Considering the most prevalent amino acids for each position.

**Figure 5:**
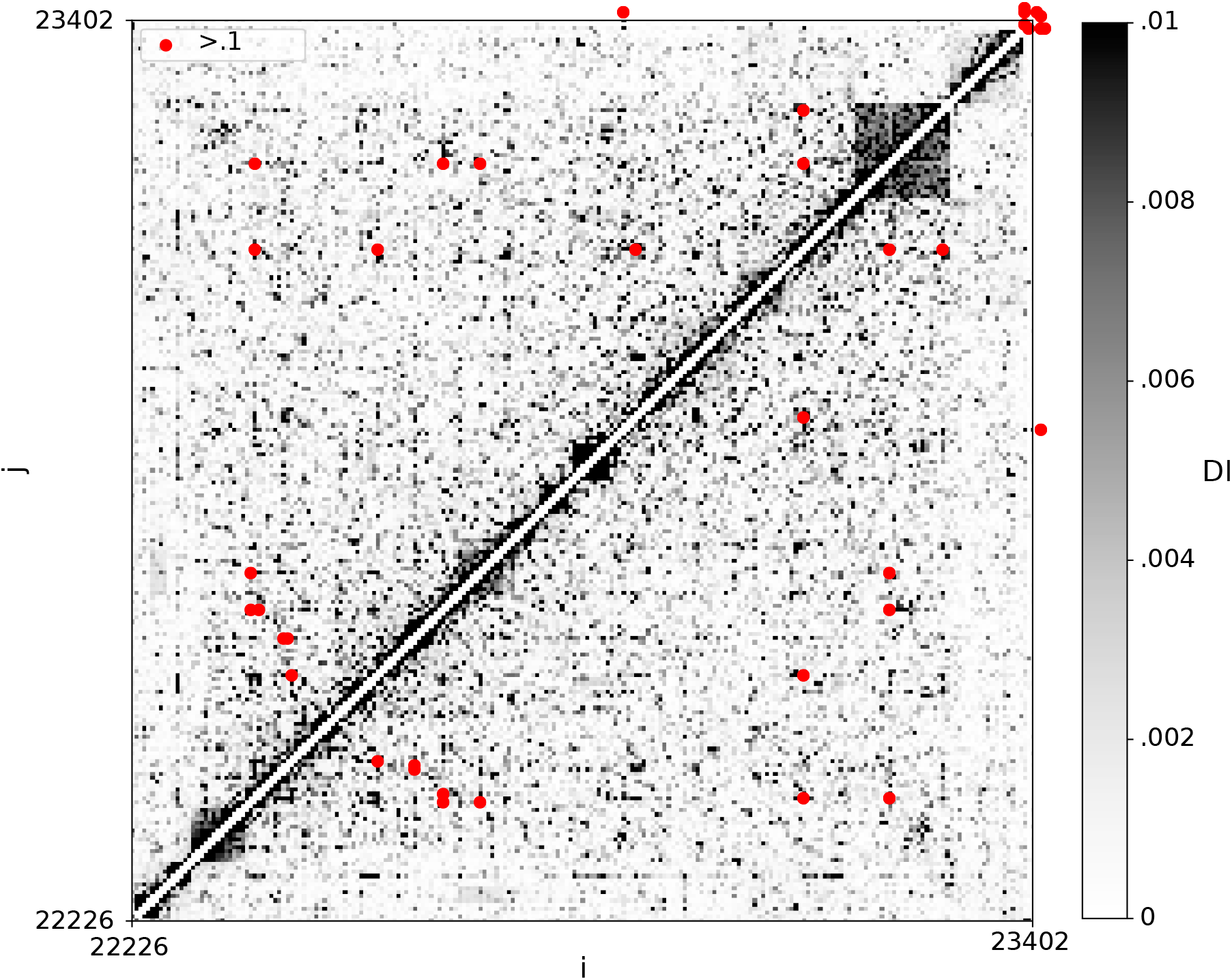
S Interaction Map.

#### 2.2.3 ORF3a

The ORF3a gene region encodes a unique membrane protein with a 3-membrane structure and it is essential for the pathogenesis of the disease [Lu et al., 2010, Issa et al., 2020].

The interaction map of ORF3a expresses minimal incidence so it is not shown. However, the interactions in ORF3a are still important to consider. The region itself shows little variation (only 5 pairs have DI*>* 0.1) with the exception of position 25563 as seen in previous work [Uğurel et al., 2020]. Regardless, in Table 7 we are able to show that 25563 interacts with several other regions including ORF1ab and S with the strongest coevolution occurring with position 1059 in the ORF1ab region. We consider the encoded amino acids for positions coevolving in ORF3a in Tables 8 and 9. In Table 9 we see a new case in the AA-pair distribution with position pairs of 25563 and both 22992 and 25429. In both these pairs, the majority of the AA pair count is the primary AA for each position. However, the second most prevalent pairing is in the primary-secondary AA couples for the position. While a high count on the matrix diagonal represents the prevalence of two AA pairs, the distribution in these pairs shows more variety with 3 significant pairings expressed in the position interaction.

**Table 7:**
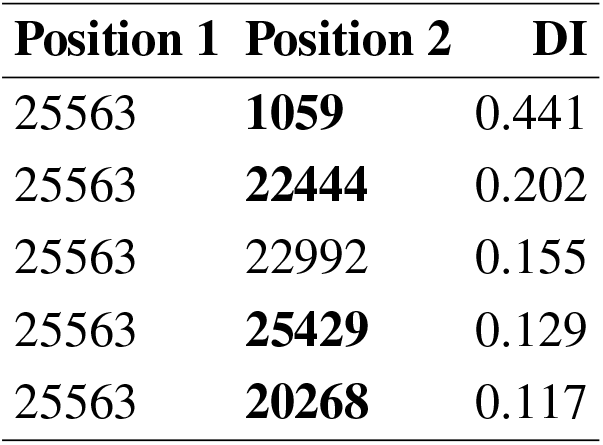
Top 5 ORF3a DI pairs. Listing the strongest DI pair positions (bolded positions not in ORF3a)

| Position 1 | Position 2 | DI |
| --- | --- | --- |
| 25563 | **1059** | 0.441 |
| 25563 | **22444** | 0.202 |
| 25563 | 22992 | 0.155 |
| 25563 | **25429** | 0.129 |
| 25563 | **20268** | 0.117 |

**Table 8:** ORF3a Single Site Amino Acid Frequencies. We consider the distributions of resulting amino acids from the nucleotide positions which had the highest DI in ORF3a

|  | Single Site<br>Freqs. |
| --- | --- |
| 25563 | Q:.783,H:.215 |
| 1059 | T:.832,I:.164 |
| 22444 | D:.891,X:.109 |
| 22992 | S:.914,N:.046,X:.039 |
| 25429 | V:.893,L:.105 |
| 20268 | L:.961,X:.039 |

**Table 9:**
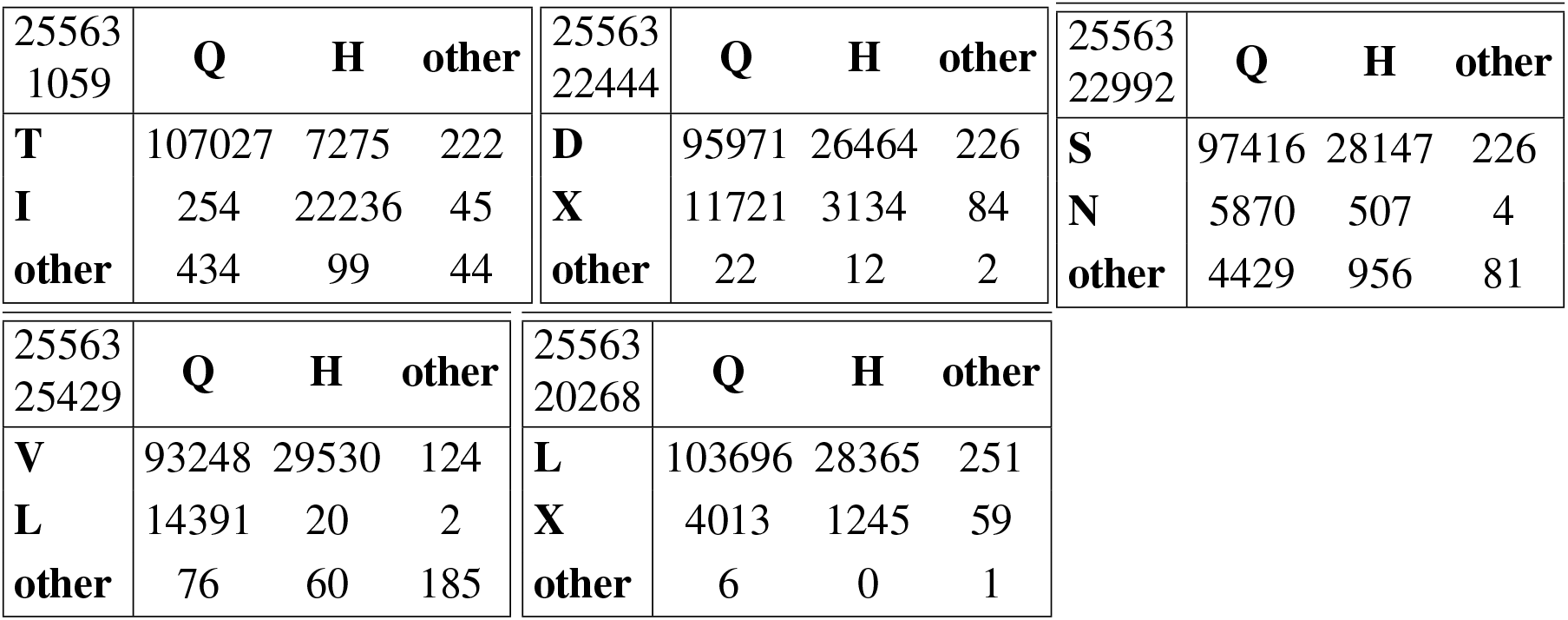
ORF3a AA Pair Counts. Encoded amino acid counts for high ranked DI positions in ORF3a.Considering the most prevalent amino acids for each position.

|  |  |  |  |  |  |  |  |  |  |  |  |
| --- | --- | --- | --- | --- | --- | --- | --- | --- | --- | --- | --- |
| 25563<br>1059 | **Q** | **H** | **other** | 25563<br>22444 | **Q** | **H** | **other** | 25563<br>22992 | **Q** | **H** | **other** |
| **T** | 107027 | 7275 | 222 | **D** | 95971 | 26464 | 226 | **S** | 97416 | 28147 | 226 |
| **I** | 254 | 22236 | 45 | **X** | 11721 | 3134 | 84 | **N** | 5870 | 507 | 4 |
| **other** | 434 | 99 | 44 | **other** | 22 | 12 | 2 | **other** | 4429 | 956 | 81 |

|  |  |  |  |  |  |  |  |
| --- | --- | --- | --- | --- | --- | --- | --- |
| 25563<br>25429 | **Q** | **H** | **other** | 25563<br>20268 | **Q** | **H** | **other** |
| **V** | 93248 | 29530 | 124 | **L** | 103696 | 28365 | 251 |
| **L** | 14391 | 20 | 2 | **X** | 4013 | 1245 | 59 |
| **other** | 76 | 60 | 185 | **other** | 6 | 0 | 1 |

#### 2.2.4 ORF7ab

ORF7ab contains a viral antagonist of host restriction factor BST-2/Tetherin and induces apoptosis [Holland et al., 2020]. We present the interaction map for both ORF7a and ORF7b separately in Figures 6 and 7. We also present the top 10 DI pairs for each region in Table 10.

**Table 10:** Top 10 ORF7ab DI pairs. Listing the strongest DI pair positions

| Position 1 | Position 2 | DI |
| --- | --- | --- |
| 27801 | 27803 | 0.365 |
| 27688 | 27792 | 0.333 |
| 27698 | 27700 | 0.299 |
| 27579 | 27581 | 0.295 |
| 27803 | 27805 | 0.277 |
| 27794 | 27796 | 0.268 |
| 27804 | 27806 | 0.248 |
| 27805 | 27807 | 0.227 |
| 27758 | 27761 | 0.209 |
| 27688 | 27752 | 0.188 |

**Figure 6:**
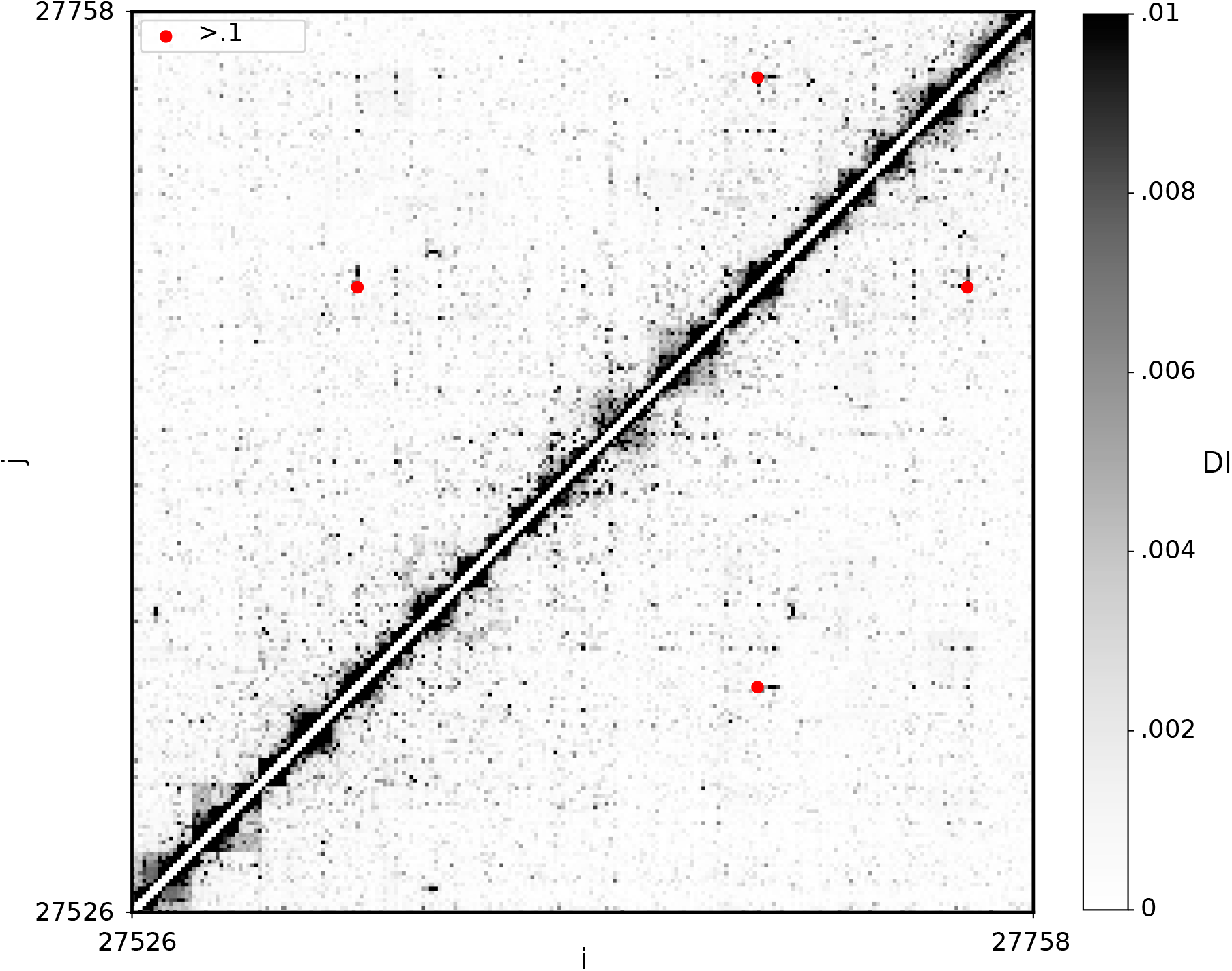
ORF7a Interaction Map.

**Figure 7:**
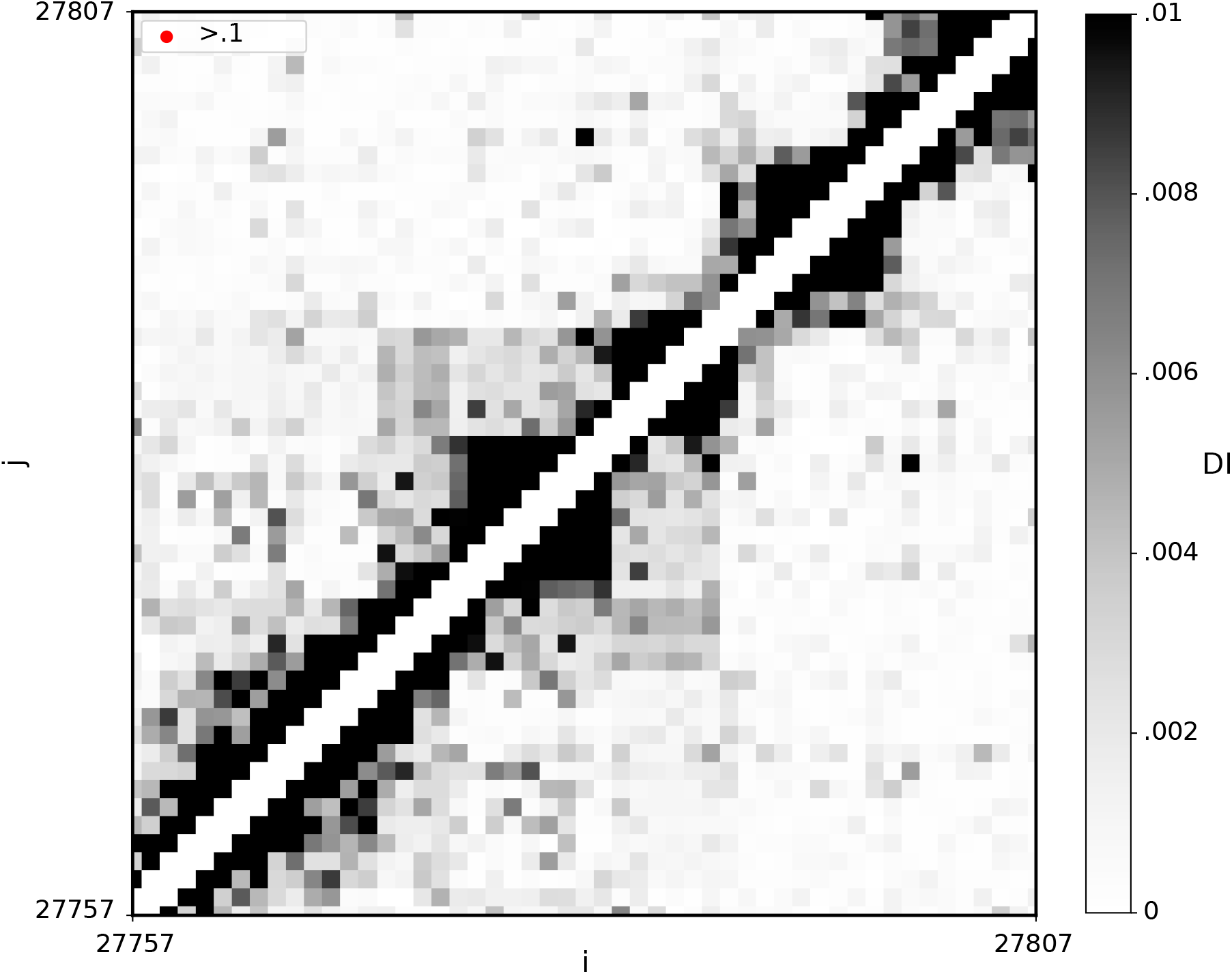
ORF7b Interaction Map.

It is important to note that there was little significant coevolution between positions in ORF7ab and other regions of the genome. We will only present the AA single site frequencies, in Table 11 for this region because of the single AA dominance at each position considered. In addition to this strong dominance (≥ 90%) of the primary AA at each position, the secondary AA at all positions was undefined (X).

**Table 11:** Single Site Amino Acid Frequencies. We consider the distributions of resulting amino acids from the nucleotide positions which had the highest DI in ORF7ab

|  | Single Site<br>Freqs. |
| --- | --- |
| 27801 | F:.934,X:.065 |
| 27803 | F:.934,X:.065 |
| 27792 | F:.929,X:.071 |
| 27688 | P:.904,X:.096 |
| 27698 | L:.904,X:.1 |
| 27700 | I:.903,X:.1 |
| 27579 | I:.902, X:.097 |
| 27581 | F:.901,X:.099 |
| 27805 | L:.941,X:.058 |
| 27794 | F:.929,X:.071 |
| 27696 | F:.931,X:.069 |
| 27804 | L:.941,X:.058 |
| 27806 | L:.941,X:.058 |
| 27807 | L:.988,X:.011 |
| 27761 | I:.902,X:.097 |
| 27752 | T:.902,X:.098 |

#### 2.2.5 Nucleocapsid

We conclude our complete genome analysis with a brief description of our findings of coevolution in the Nucleocapsid (N) encoding region. The N protein plays varied roles in the regulation of the infected cell metabolism and packaging of the viral genome. Therefore, the protein plays an important role in both replication and transcription [Kang et al., 2020]. The N region contains an specific nucleotide variation in the triplet 28881, 28882, and 28883 discussed in previous work [Uğurel et al., 2020].This triplet, appropriately, presents the only DI≥ 0.1 in the region, as presented in Table 12.

**Table 12:** Nucleocapsid (N) DI pair. The single pairing ≥ 0.1 is also one of the strongest DI in the genome.

| Position 1 | Position 2 | DI |
| --- | --- | --- |
| 28881 | 28883 | 0.801 |

### 2.3 Clades

When investigating genetic variance, it is useful to stratify available data to understand and analyse genomic diversity. Analysis of genetic variance plays a crucial role in expanding knowledge and developing prevention strategies. Previous work has developed phylogenetic trees and divided the SARS-CoV-2 genome both genomically and geographically into clades [Mercatelli & Giorgi, 2020]. We extend this analysis to see how such stratification affects the virus’s genome-wide covariation. Before presenting our results on clades, it is important to note that our initial results yielded many previously defined clade determinants [Mercatelli & Giorgi, 2020]. These determinant nucleotide (NT) positions are bolded in Table 13. In the following sections we apply our method to these clades and investigate the resulting change in the coevolution of NT positions across the genome.

**Table 13:** Clade Determinants in Genome-Wide Analysis. We outline the clade determinants from DI presented in Section 2.2 (bolded)

| Clade Determinants Present in DI |  |  |  |  |
| --- | --- | --- | --- | --- |
| Position 1 | Position 2 | DI | Region | Clade |
| **14408** | **3037** | 0.456 | ORF1ab | G |
| 1059 | **25563** | 0.441 | ORF1ab | GH |
| 14805 | 26144 | 0.380 | ORF1ab | V |
| **14408** | 23404 | 0.340 | ORF1ab | G |
| **3037** | **23403** | 0.335 | ORF1ab | G, GR |
| **25563** | 1059 | 0.441 | ORF3a | G |
| **25563** | 22444 | 0.202 | ORF3a | G |
| **25563** | 22992 | 0.155 | ORF3a | G |
| **25563** | 25429 | 0.129 | ORF3a | G |
| **25563** | 20268 | 0.117 | ORF3a | G |
| **28881** | **28883** | 0.801 | N | GR |

#### 2.3.1 G Clade

We begin our clade analysis with the largest existing clade. The G clade is stratified by the most common set of events, a quadruplet of mutations: C241T, C3037T, C14408T, A23403G [Mercatelli & Giorgi, 2020]. Extracting genome sequences with these features we re-apply ER, resulting in the interaction map in Figure 8. Comparing the full interaction map (Figure 3) and the G clade interaction map (Figure 8), we see a drastic change in incidence.

**Figure 8:**
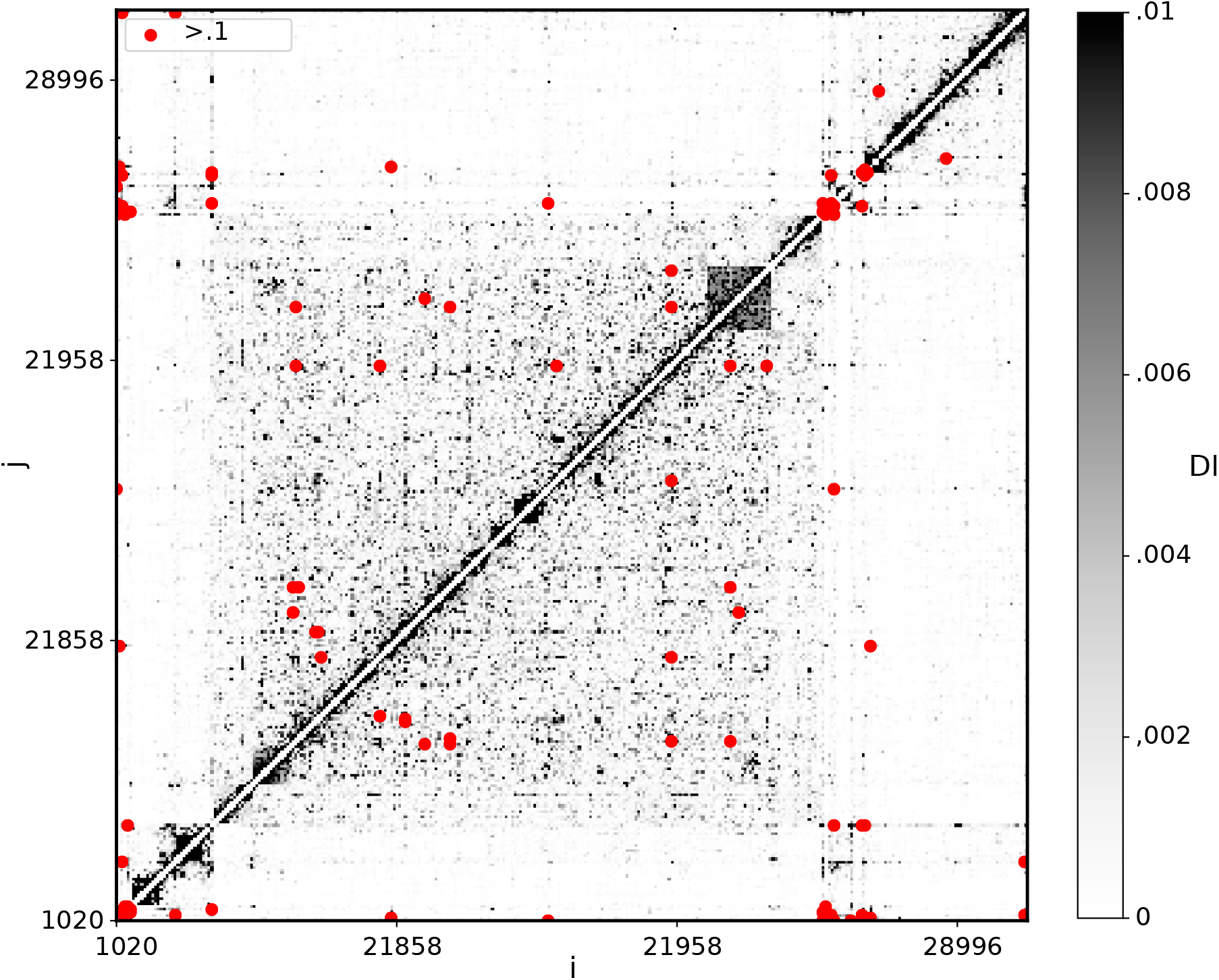
G Clade Full Genome Interaction Map.

Both genome sequence sets have the same curation applied, with pre-processing removing genome positions which were ≥ 95% conserved. However, within the G clade we see a severely decreased incidence in ORF1ab, with positions from NT 19300 to 19500 no longer showing sufficient variation. This change in incidence is expressed in Figure 9 as a decrease in cardinality of the interaction map from the full genome set to the G clade genome set.

**Figure 9:**
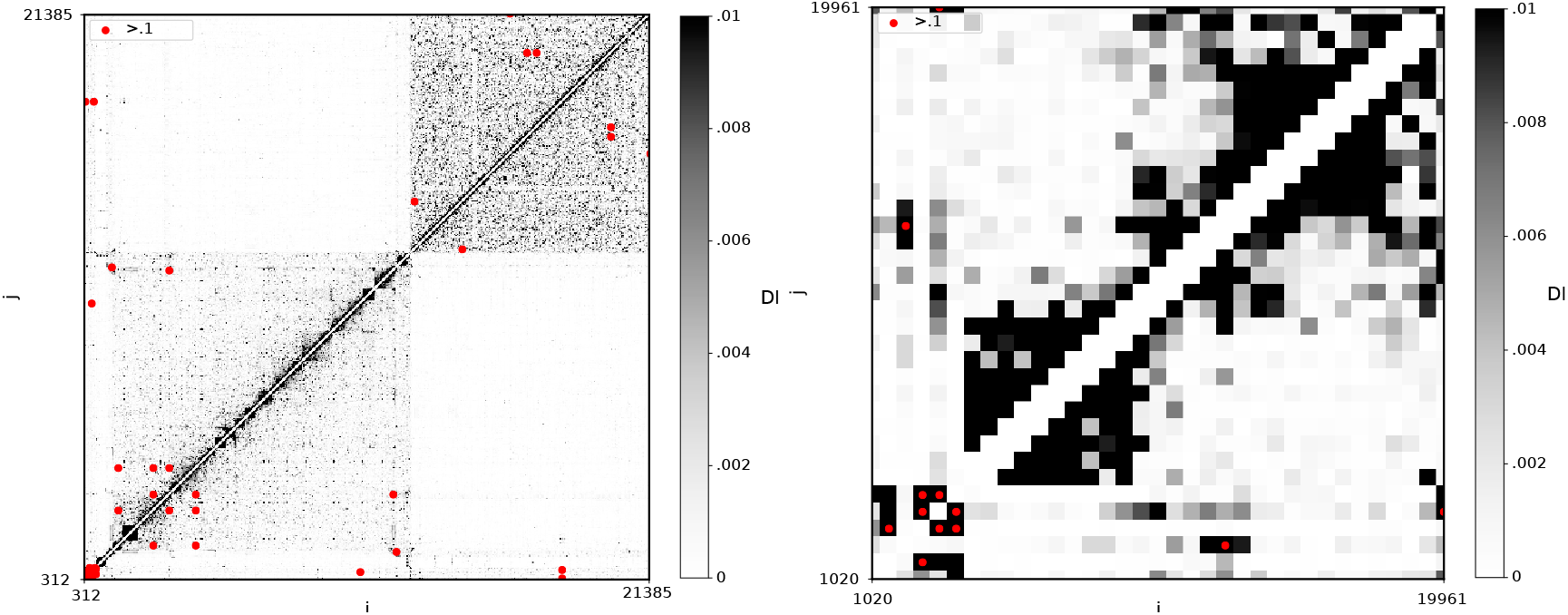
ORF1ab Interaction Maps. Showing the interaction map for the ORF1ab region with both the full genome sequence set (left), as well as the G clade genome sequence set (right)

The incidence change in the S encoding region is much less severe. The NT range from 2200 to 24000 appears mostly unchanged, showing the same pattern of significant DI (*DI >* 0.1, red dots).In addition to the changed incidence of ORF1ab and S. While variation of the different encoding regions differs in ORF1ab and S, there are alterations in the top ranked DI positions throughtout both regions.

First consider Table 14, which gives the top ranked DI pairs in ORF1ab for the full genome sequence set and the G clade genome sequence set side by side. None of the top ranked DI pairs from the full genome analysis remain in the top ranked pairs for the G clade. This is not due to an overall loss in information, as the DI magnitude remains in the same region (DI∈ [0.2, 0.4]). Most of the removed pairs were G clade determinants, therefore these positions are fixed in the G clade, specifically positions 14408, 3037, 23403 (and proximal positions). By fixing these positions we effectively removed the variation of positions represented in 6 of the top 10 pairs. However, it seems that the connection between ORF1ab and other encoding regions, specifically the S region, was retained. Consider the NT position 23403 from the S encoding region, which was fixed in the G clade genome sequence set. During the stratification of the G clade genome sequence set, the variation at 23403 (likely 23401 and 23404 as well) was removed. This small NT group in region S accounted for 4 of the top 10 DI pairs for ORF1ab. However, while 23403 and its corresponding positions no longer coevolved with ORF1ab, the NT position 22870 expressed a very strong connection with ORF1ab in the G clade.

**Table 14:**
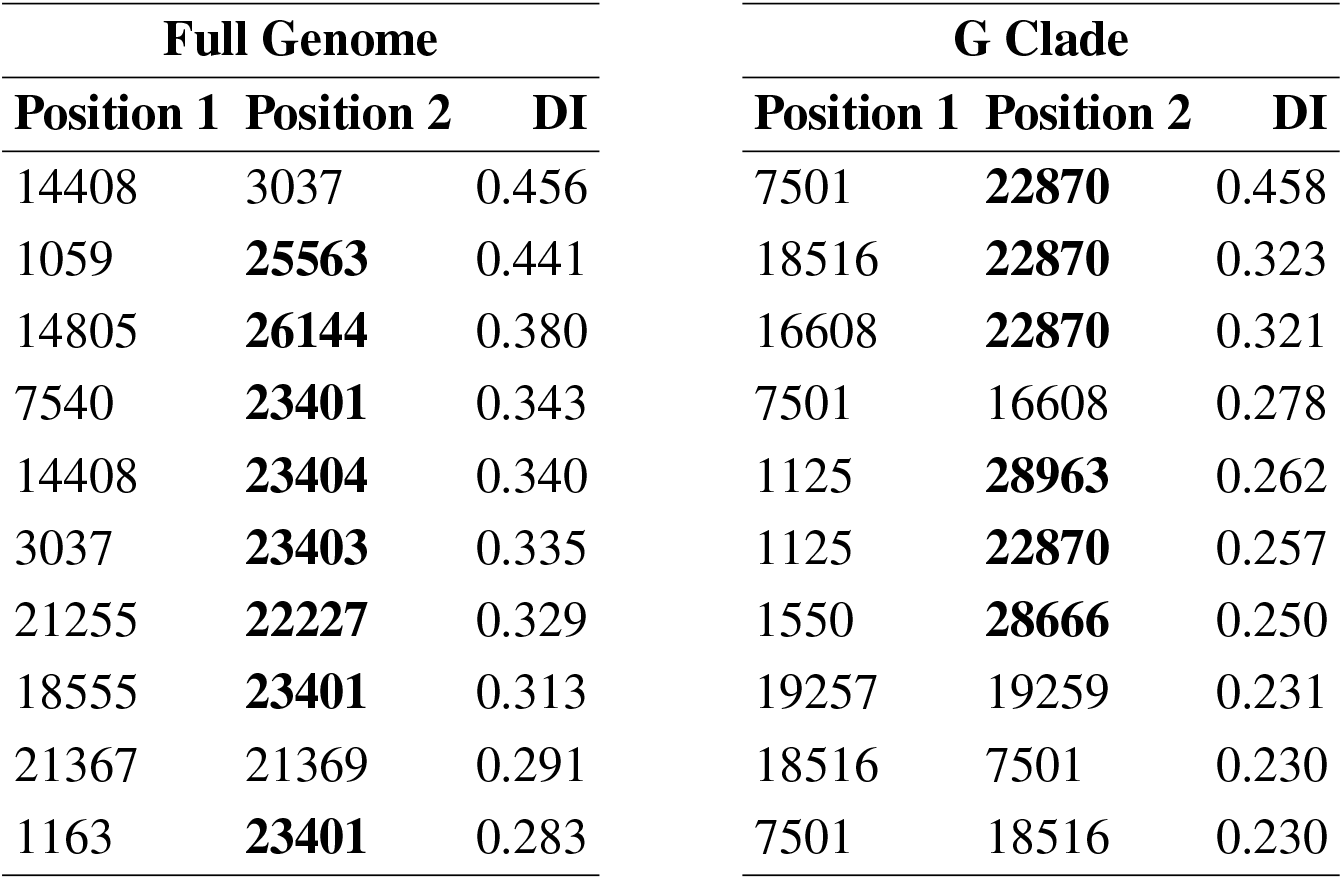
ORF1ab top ranked DI pairs. We show the 10 top ranked DI pairs for ORF1ab generated from the full genome sequence set and the G clade genome sequence set

This trend continues in the S encoding region. Table 15 shows the top ranked DI pairs in the S encoding region for the full genome sequence set and the G clade genome sequence set. As before, none of the top ranked DI pairs from the full set survive the stratification which creates the G clade genome sequence set.

**Table 15:** S top ranked DI pairs. We show the 10 top ranked DI pairs for S generated from the full genome sequence set and the G clade genome sequence set

| Full Genome |  |  | G Clade |  |  |
| --- | --- | --- | --- | --- | --- |
| Position 1 | Position 2 | DI | Position 1 | Position 2 | DI |
| 22363 | 22384 | 0.915 | 21832 | 21853 | 0.803 |
| 23401 | **7540** | 0.343 | 22870 | **7501** | 0.458 |
| 23401 | **18555** | 0.313 | 22870 | **18516** | 0.323 |
| 22497 | 22495 | 0.301 | 22870 | **16608** | 0.321 |
| 22353 | 22355 | 0.292 | 21822 | 21824 | 0.265 |
| 23401 | **1163** | 0.283 | 21966 | 21964 | 0.264 |
| 23401 | **16647** | 0.275 | 21964 | 21966 | 0.263 |
| 22334 | 22336 | 0.263 | 22870 | **1125** | 0.257 |
| 22539 | 22541 | 0.245 | 21995 | 21993 | 0.244 |
| 22526 | 22524 | 0.239 | 22008 | 22010 | 0.236 |
| 22354 | 22488 | 0.218 | 21803 | 21805 | 0.234 |

#### 2.3.2 Other Clades

We conclude our analysis with an overview of the remaining clades presented in previous work: GR, GH, S, and V clades [Mercatelli & Giorgi, 2020]. Specifically we present the interaction maps for the different clades (Figures 10-13).

**Figure 10:**
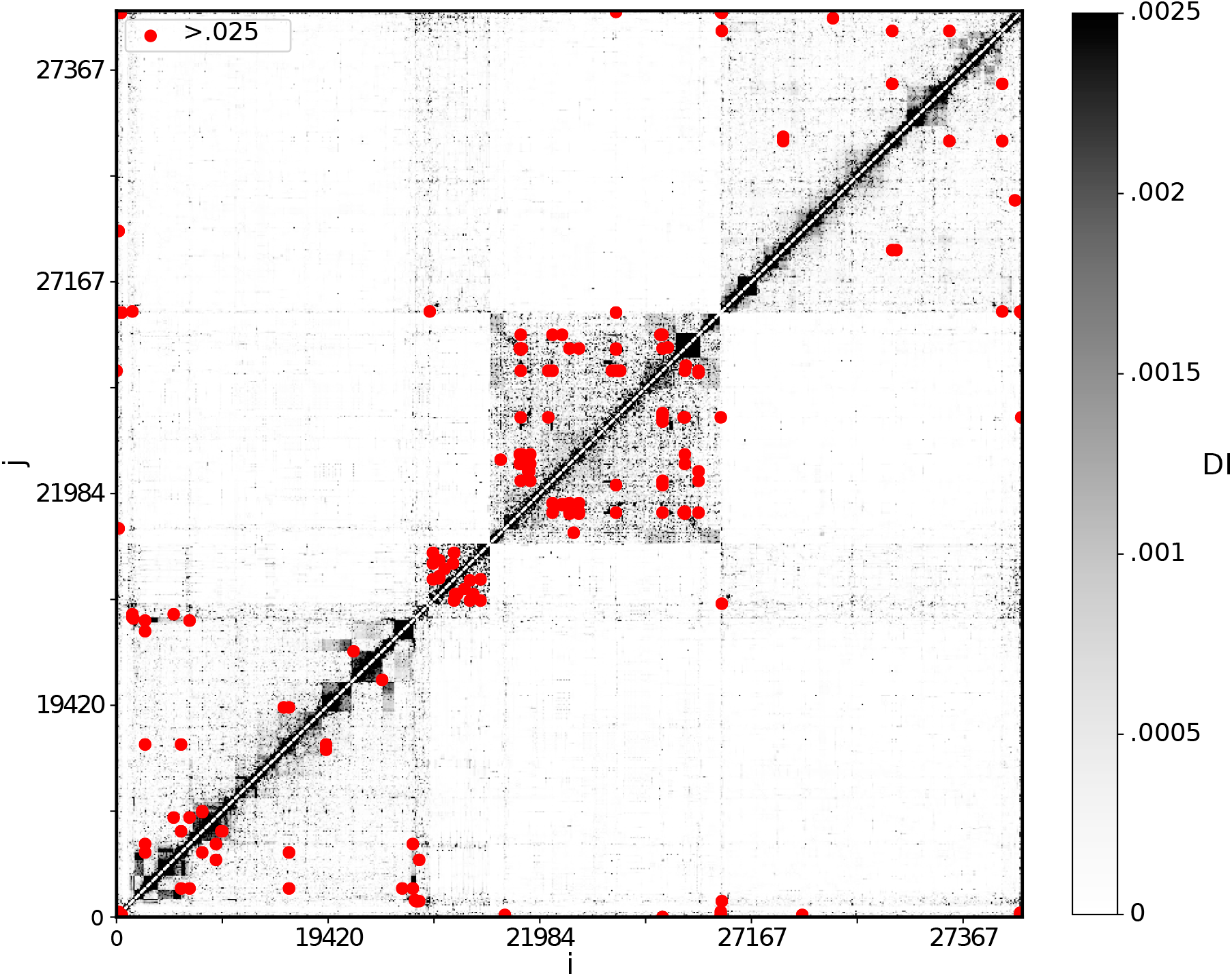
GH Clade Interaction Map. Showing the interaction map for the GH clade genome sequence set

**Figure 11:**
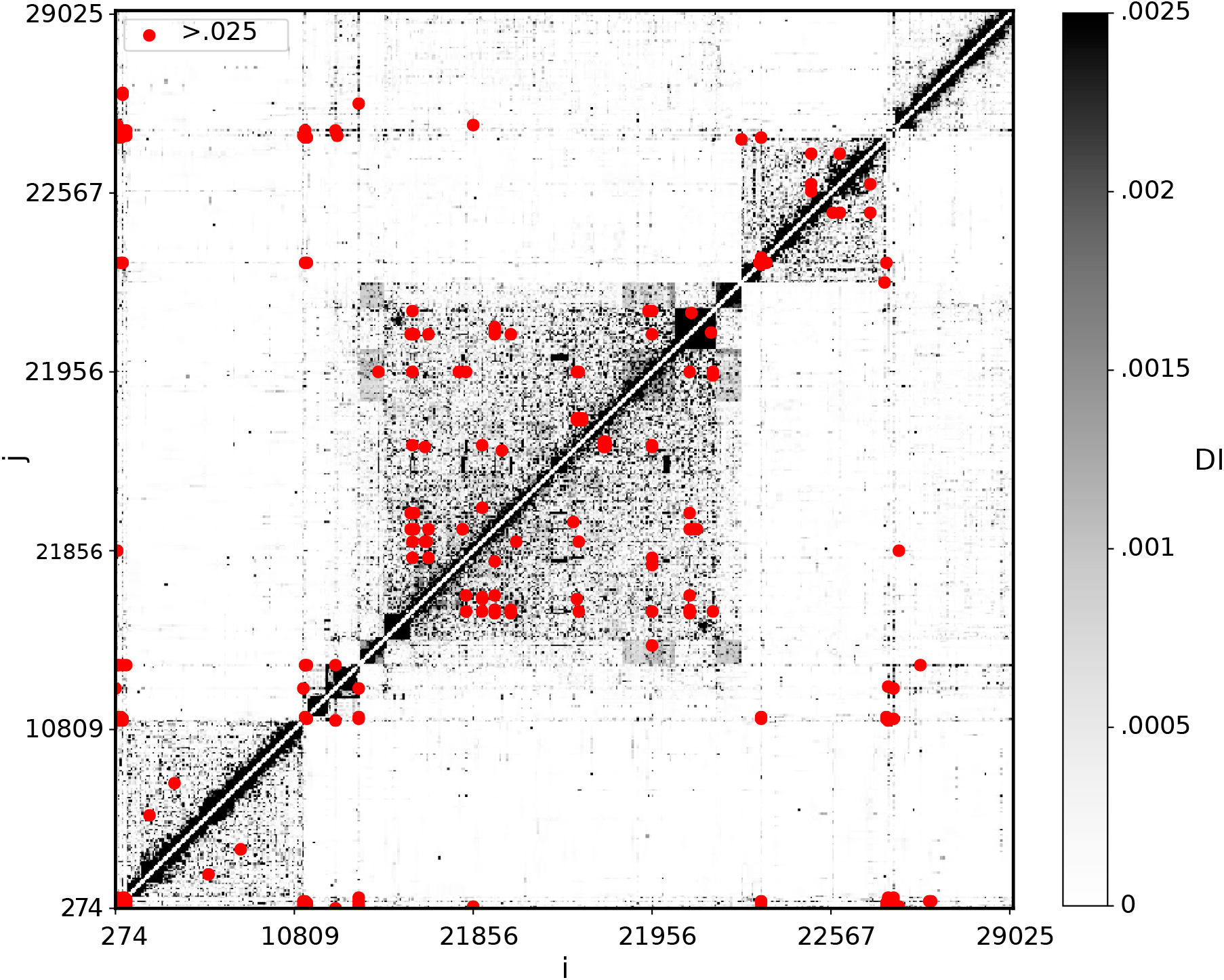
GR Clade Interaction Map. Showing the interaction map for the GR clade genome sequence set

**Figure 12:**
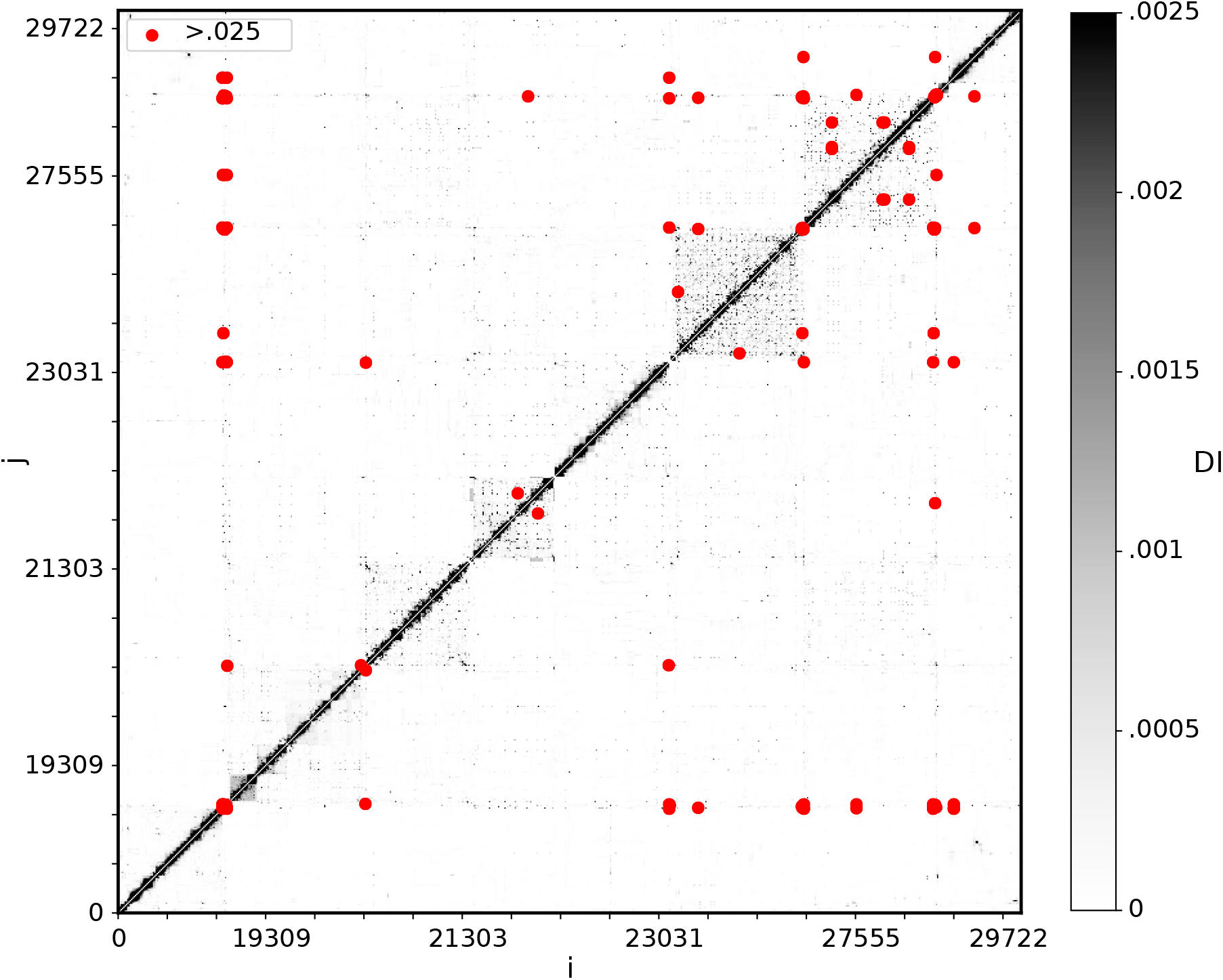
S Clade Interaction Map. Showing the interaction map for the S clade genome sequence set

**Figure 13:**
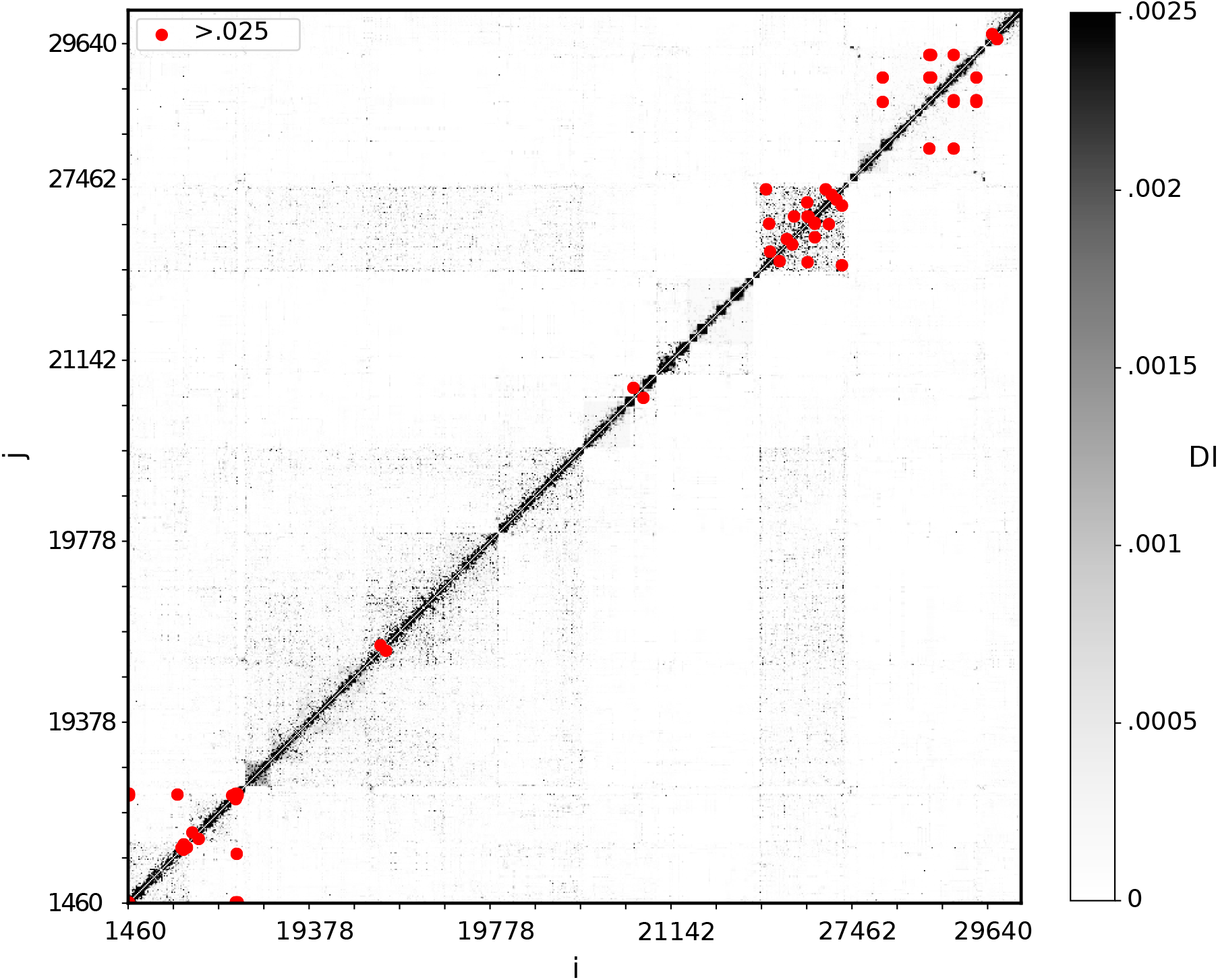
V Clade Interaction Map. Showing the interaction map for the V clade genome sequence set

With the given number of sequences in the smaller clades, the sample size limits the inference. This is evident when considering the shift in DI threshold in the Figures 10-13 with significant DI changing 0.1 −→ 0.025. Due to the smaller sample sizes in these clades, we forego further analysis as the inference would be unreliable.

## 3 Discussion

In this work we have presented a novel analysis of the SARS CoV-2 genome by finding genome interactions both within and across encoding regions. In our analysis of the SARS CoV-2 genome we have presented several perspectives on nucleotide interactions in the genome. These interactions showed both proximal interactions within individual encoding regions as well as distal interactions between different encoding regions throughout the genome. We inferred nucleotide position interactions for the entire genome as well as the separate encoding regions. Particular attention was given to the ORF1ab and S regions, which demonstrated the highest variability in the given dataset. We were able to draw analogous conclusions from previous work by inferring the the most common variations previously reported [Uğurel et al., 2020]. Additionally, our genome-wide interaction maps expressed determinant positions of all clades available at the time our data was acquired [Mercatelli & Giorgi, 2020]. Generating interaction maps of individual clades showed clade-specific coevolution of nucleotide positions.

We further considered the level of variability, both within regions of the full genome data set and for different clades. This was accomplished by varying the threshold for conserved columns while considering the retained column incidence. This relationship shows nucleotide variability is different both between encoding regions of the full genome and between different clades. Region-specific incidences are not consistent between clades, with individual regions expressing different variability in different clades. Comparison of region-specific incidence can also give intuition on the level of variability or specific regions within specific clades.

Future extensions of this analysis provide several avenues of investigations. First, as the database of SARS CoV-2 genomes grows, the incidence and overall variability will increase, yielding further insights into genome interactions. Additionally, the availability of data over longer time periods will allow for chronological compartmentalization of genome data sets and interaction maps can be compared across the temporal evolution of the virus. Second, this analysis can also be applied to diseases for which there is more data available as the importance of genome interactions is not SARS-CoV-2 specific.

## 4 Materials and Methods

### 4.1 Expectation Reflection

To infer position pair couplings we first utilize a data-driven method, Expectation Reflection (ER), which outperforms current methods in inferring network interactions between binary variables, especially for small sample sizes[Hoang et al., 2019b]. Here we outline the method and how it is applied to infer connections between genome positions. We begin with a given genetic sequence,

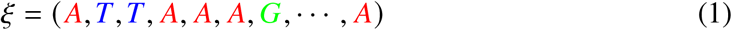

In order to translate this into a binary variable sequence required for Expectation Reflection we can use a OneHot transformation [Pedregosa et al., 2011]. This transformation converts a nucleotide into a binary representation,

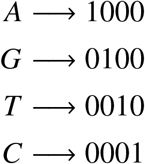

which allows us to convert the genome sequence set into a binary sequence set. Therefore the previous sequence in Equation 1 becomes *σ* = OneHot(ξ),

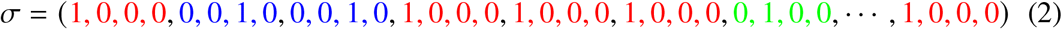

Now that we have established the conversion of a set of genetic sequences into a set of binary variables we can continue to the application of Expectation Reflection. Given a binary variable *σ* (*t*) representing the *t*^*th*^ sequence such that with *t* ∈ [1, *N* = 137686] sequences, or states, which the SARS-CoV-2 virus genome can take. An individual sequence has the form *σ* = (*σ*_1_, *σ*_2_, …, *σ*_*NT*_) where *NT* is the number of binary variables representing the nucleotides such that with the 29903 length genome *NT* = 29903 *times*4. We can then assume that given a current sequence *σ* (*t*), an individual position in a future sequence changes stochastically according to the following conditional probability,

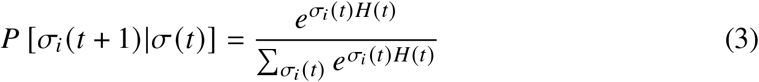

where the local field, *H*_*i*_ (*t*),

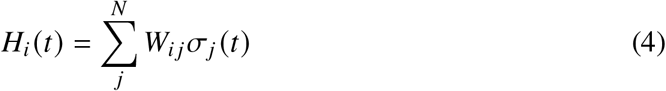

where *W*_*i j*_ represents the connection between positions *i* and *j*. *H*_*i*_ is a function of the current sequence state *σ* (*t*)_*i*_ and expresses the influence of a given sequence position *σ*_*j*_ on the future state of sequence position *σ*_*i*_. This conditional relationship allows us to iteratively search for the ideal *W* using all *N* available sequences. The method, and resulting algorithm, is discussed in further detail in previous work [Hoang et al., 2019a, Hoang et al., 2019b]

With the resulting coupling matrix we can calculate the Direct Information (DI) between all position pairs using Direct Coupling Analysis, which has been widely used to infer such interactions in previous work. [Weigt et al., 2009, Morcos et al., 2011, Ekeberg et al., 2014, De Leonardis et al., 2015].

### 4.2 Genome Data: Acquisition and Alignment

The data used for both the full genome-wide covariation and the clade-specific genomewide covariation was acquired from GISAID [Shu & McCauley, 2017]. We downloaded all 137636 available complete SARS-CoV-2 sequences on October 19, 2020. The resulting sequences were then aligned on the Biowulf Linux cluster at the National Institutes of Health, Bethesda, MD. MAFFT [Katoh et al., 2005] was used to align the sequences.

## 5 Acknowledgements

The authors would like to thank Prakriti Mudvari and Eli Boritz from the National Institute of Allergy and Infectious Diseases (NIAID) for help with data acquisition. Additionally, the authors would like to thank the HPC staff, Wolfgang Resch and Gennady Denisov for computational support using Biowulf. This work utilized the computational resources of the NIH HPC Biowulf cluster (http://hpc.nih.gov). This work was supported by the Intramural Research Program of the National Institute of Diabetes and Digestive and Kidney Diseases.

## References

Briguglio, I., Piras, S., Corona, P., & Carta, A. (2011). Inhibition of RNA helicases of ssRNA+ virus belonging to Flaviviridae, Coronaviridae and Picornaviridae families. International journal of medicinal chemistry, 2011.

Dahirel, V., et al. (2011). Coordinate linkage of HIV evolution reveals regions of immunological vulnerability. Proceedings of the National Academy of Sciences, 108(28), 11530–11535, https://doi.org/10.1073/pnas.1105315108. https://www.pnas.org/content/108/28/11530.

De Leonardis, E., Lutz, B., Ratz, S., Cocco, S., Monasson, R., Schug, A., & Weigt, M. (2015). Direct-Coupling Analysis of nucleotide coevolution facilitates RNA secondary and tertiary structure prediction. Nucleic acids research, 43(21), 10444–10455.

Ekeberg, M., Hartonen, T., & Aurell, E. (2014). Fast pseudolikelihood maximization for direct-coupling analysis of protein structure from many homologous amino-acid sequences. Journal of Computational Physics, 276, 341–356.

Gallagher, T. M. & Buchmeier, M. J. (2001). Coronavirus spike proteins in viral entry and pathogenesis. Virology, 279(2), 371–374.

Gorbalenya, A., et al. (2020). The species severe acute respiratory syndrome related coronavirus: classifying 2019-nCoV and naming it SARS-CoV-2. Nat Microbiol 5: 536–544.

Hoang, D.-T., Jo, J., & Periwal, V. (2019a). Data-driven inference of hidden nodes in networks. Physical Review E, 99(4), 042114.

Hoang, D.-T., Song, J., Periwal, V., & Jo, J. (2019b). Network inference in stochastic systems from neurons to currencies: Improved performance at small sample size. Physical Review E, 99(2), 023311.

Holland, L. A., et al. (2020). An 81-Nucleotide Deletion in SARS-CoV-2 ORF7a Identified from Sentinel Surveillance in Arizona (January to March 2020). Journal of Virology, 94(14), https://doi.org/10.1128/JVI.00711-20. https://jvi.asm.org/content/94/14/e00711-20.

Issa, E., Merhi, G., Panossian, B., Salloum, T., & Tokajian, S. (2020). SARS-CoV-2 and ORF3a: Nonsynonymous Mutations, Functional Domains, and Viral Pathogenesis. Msystems, 5(3).

John Hopkins University, J. (2020). Coronavirus Resource Center. https://coronavirus.jhu.edu/.

Kang, S., et al. (2020). Crystal structure of SARS-CoV-2 nucleocapsid protein RNA binding domain reveals potential unique drug targeting sites. Acta Pharmaceutica Sinica B.

Katoh, K., Kuma, K.-i., Toh, H., & Miyata, T. (2005). MAFFT version 5: improvement in accuracy of multiple sequence alignment. Nucleic acids research, 33(2), 511–518.

Kwong, A. D., Rao, B. G., & Jeang, K.-T. (2005). Viral and cellular RNA helicases as antiviral targets. Nature reviews Drug discovery, 4(10), 845–853.

Lu, W., Xu, K., & Sun, B. (2010). SARS accessory proteins ORF3a and 9b and their functional analysis. In Molecular Biology of the SARS-Coronavirus (pp. 167–175). Springer.

Mercatelli, D. & Giorgi, F. M. (2020). Geographic and Genomic Distribution of SARS-CoV-2 Mutations. Frontiers in Microbiology, 11, 1800, https://doi.org/10.3389/fmicb.2020.01800. https://www.frontiersin.org/article/10.3389/fmicb.2020.01800.

Morcos, F., et al. (2011). Direct-coupling analysis of residue coevolution captures native contacts across many protein families. Proceedings of the National Academy of Sciences, 108(49), E1293–E1301.

NCBI (2020). Accession No: NC_045512.2. Severe acute respiratory syndrome coronavirus 2 isolate Wuhan-Hu-1. https://www.ncbi.nlm.nih.gov/nuccore/NC_045512.

Pedregosa, F., et al. (2011). Scikit-learn: Machine Learning in Python. Journal of Machine Learning Research, 12, 2825–2830.

Shu, Y. & McCauley, J. (2017). GISAID: Global initiative on sharing all influenza data–from vision to reality. Eurosurveillance, 22(13), 30494.

Song, Z., et al. (2019). From SARS to MERS, thrusting coronaviruses into the spotlight. Viruses, 11(1), 59.

Tai, W., et al. (2020). Characterization of the receptor-binding domain (RBD) of 2019 novel coronavirus: implication for development of RBD protein as a viral attachment inhibitor and vaccine. Cellular & molecular immunology, 17(6), 613–620.

Uğurel, O. M., Ata, O., & Balik, D. (2020). An updated analysis of variations in SARS-CoV-2 genome. Turkish Journal of Biology, 44(SI-1), 157–167.

Weigt, M., White, R. A., Szurmant, H., Hoch, J. A., & Hwa, T. (2009). Identification of direct residue contacts in protein–protein interaction by message passing. Proceedings of the National Academy of Sciences, 106(1), 67–72.

World Health Organization, W. (2020). Coronavirus disease (COVID-2019) situation re-ports. http://web.archive.org/web/20080207010024/ http://www.808multimedia.com/winnt/kernel.htm.

Wu, A., et al. (2020). Genome composition and divergence of the novel coronavirus (2019-nCoV) originating in China. Cell host & microbe.

Zhou, Y., Hou, Y., Shen, J., Huang, Y., Martin, W., & Cheng, F. (2020). Network-based drug repurposing for novel coronavirus 2019-nCoV/SARS-CoV-2. Cell discovery, 6(1), 1–18.

Zhu, N., et al. (2020). A novel coronavirus from patients with pneumonia in China, 2019. New England Journal of Medicine.

